# Droplet scRNA-seq is not zero-inflated

**DOI:** 10.1101/582064

**Authors:** Valentine Svensson

## Abstract

Potential users of single cell RNA-sequencing often encounter a choice between high-throughput droplet based methods and high sensitivity plate based methods. In particular there is a widespread belief that single-cell RNA-sequencing will often fail to generate measurements for particular gene, cell pairs due to molecular inefficiencies, causing data to have an overabundance of zero-values. Investigation of published data of technical controls in droplet based single cell RNA-seq experiments demonstrates the number of zeros in the data is consistent with count statistics, indicating that over-abundances of zero-values in biological data are likely due to biological variation as opposed to technical shortcomings.

## Background

As single cell RNA-sequencing (scRNA-seq) started to gain popularity users expressed concern about an unexpected number of zero values among gene expression measures. That is, for any given gene many cells had not detected the expression, even if the level was relatively high in other cells (Bacher and Kendziorski, 2016; Haque et al., 2017; Vallejos et al., 2017). Often this is considered distinct from “true zeros”, which are zeros due to the gene not being expressed.

For context, it is worth tracing the history of handling zeros in scRNA-seq data: It was reported in 2014 that genes in scRNA-seq data had a higher number of zeros when comparing two samples on a logged reads-per-million scale than was typically observed when comparing two bulk RNA-seq samples (Kharchenko et al., 2014). This was related to studies in single cell reverse-transcriptase qPCR (sc-qPCR) where data was considered was zero-inflated but otherwise log-normal (McDavid et al., 2013). To appropriately solve the task of finding shifts in mean between conditions, it was in both cases concluded that a zero-inflation model would be needed. The sc-qPCR method used a zero-inflated log-normal model, while the scRNA-seq method SCDE used a zero-inflated negative binomial approach.

Interestingly, without much comment, the first version of the popular Monocle tool included special handling of zeros: the differential expression test used was a tobit model on log(FPKM) values (Trapnell et al., 2014). A tobit model is a normal (continuous) distribution which is censored on the left.

Around the same time it was reported that as part of biological variation between cells of the same type gene expression levels were stochastically bimodal (Shalek et al., 2013). This was observed when looking at expression levels of replicate cells on log(TPM + 1) scales. This was taken as contrasting with fluorescence studies of protein levels in single cells, where no such bimodality was observed (Sigal et al., 2006), and it was concluded that cells had distinct off-states from variable on-states at the RNA level.

These early studies viewed zeros differently, either they were considered a technical artefact, and in the other a biological effect. Regardless, a couple of years later a study was published which aimed to solve the problem of performing factor analysis while accounting for excessive zeros in in scRNA-seq data, called ZIFA (Pierson and Yau, 2015). In this paper it was noticed that on the log(read count) unit the probability of observing a zero was

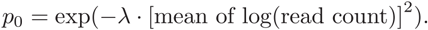

The model for the data was normal (on log-scaled data) with an added probability of zero, thus treating the data as continuous with additional zeros.

A related method CIDR looks at expression on the log(TPM) scale and aims to perform MDS on the data. To achieve this the authors construct a dissimilarity measure which is aware of zeros, but otherwise euclidean (Lin et al., 2017).

The authors describing the statistical test MAST pointed out bimodality of expression observed in scRNA-seq data, taking observational form as zeros, hypothesized it being due to bursty transcription kinetics in cells (Finak et al., 2015). Gene expression on the log2(TPM + 1) scale were modeled in a sequential fashion: first whether the gene will have a value or not, then if it is predicted to have a value, the distribution is taken as normal. Here the rate of zeros is predicted as a function of the total number of genes detected in a cell.

In a study on the design of scRNA-seq experiments researchers found a relation between the dropout rate (the fraction of cells in which the expression is zero) and log10 of mean expression on the CPM scale (as well as a log10 of the variance of expression on the CPM scale) (Tung et al., 2017).

The typical view of dropouts is that they are genes which are present (have high expression) but are still not observed in some cells. In this view the inclusion of UMI’s, which removes the effect of transcript lengths and PCR duplication, will not solve the problem of dropouts causing zero-inflation (Vallejos et al., 2017).

It has often been noted that the dropout rate relates to the read depth per cell, and since droplet scRNA-seq data has lower sequencing depth the assumption is that these are even more zero-inflated. This has led to two paths of further development.

On a statistics oriented branch of the scRNA-seq methods field, the zero-inflated negative binomial distribution have become popular as a way to directly model scRNA-seq data (Eraslan et al., 2018; Lopez et al., 2018; Risso et al., 2018).

On a computer science oriented branch of the scRNA-seq methods field, many methods have been designed to correct dropout zeros in data, with the aim of letting a user predict what the expression level of a gene in a cell would have been, had there been no zero-inflation or dropouts (Azizi et al., 2017; van Dijk et al., 2018; Gong et al., 2018; Huang et al., 2018; Li and Li, 2018; Tang et al., 2018; Zhu et al., 2016).

Closer investigations of data seem to counter the assumption that zero inflation is an inherent property of scRNA-seq data. In a study on how to perform accurate simulations for scRNA-seq data, the authors found negative binomial to be sufficient for UMI data, with little added benefit from a zero-inflation component (Vieth et al., 2017). Similarly, the authors of the method bayNorm found that zero-inflation was not necessary (Tang et al., 2018). One group proposed that if data is assumed to follow negative binomial noise, genes with excessive zeros might be good candidates for further analysis (Andrews and Hemberg, 2018).

To further investigate droplet-based high throughput single-cell RNA-seq platforms in terms of zero-inflation, this study makes use of negative control data, where absolutely no biological heterogeneity is expected. This will answer whether technical shortcomings in scRNA-seq methods produces an excess of zeros compared to expectations^1^.

## Results

In droplet-based scRNA-seq methods, negative control data can be generated by adding a solution of RNA to the fluid in microfluidic systems, making the RNA content in each droplet identical. Five such datasets have been published: one used to benchmark Drop-seq (Macosko et al., 2015); one used to benchmark InDrops (Klein et al., 2015); one used to benchmark an early version of the commercial scRNA-seq platform from 10x Genomics (Zheng et al., 2017); and two used to benchmark a later version of the commercial platform from 10x Genomics (Svensson et al., 2017). The negative control data differs in the RNA solution added (Table 1). For simplicity, each RNA species regardless of source will be referred to as a “gene” in the text. All negative control experiments aimed to make an RNA dilution that would fill each droplet with an RNA abundance similar to the content of a typical mammalian cell.

**Table 1.** Data and results. Description of the negative control data used, the fitted overdispersion values (Phi) and the results. “Variance correlation” refers to the Spearman correlation between the observed empirical variance for each gene and the variance expected given the mean and overdispersion. “Expected zeros correlation” refers to the Spearman correlation between the observed number of zeros for a gene and the expected number of zeros given the overdispersion and mean.

| Dataset | Technique | Input material | Droplets | Observed genes | Phi | Variance correlation | Expected zeros correlation |
| --- | --- | --- | --- | --- | --- | --- | --- |
| Klein et al 2015 | InDrops | K562 cell line RNA + ERCC spike-ins | 953 | 25266 | 0.04 | 1 | 1 |
| Macosko et al 2015 | Drop-seq | ERCC spike-ins | 84 | 959 | 0.12 | 0.99 | 0.98 |
| Zheng et al 2017 | GemCode | ERCC spike-ins | 1015 | 91 | 0.04 | 1 | 0.99 |
| Svensson et al 2017 (1) | Chromium (v1) | Human control brain RNA + ERCC spike-ins | 2000 | 20647 | 0.09 | 1 | 1 |
| Svensson et al 2017 (2) | Chromium (v1) | Human control brain RNA + ERCC spike-ins | 2000 | 21411 | 0.37 | 1 | 1 |

A somewhat flexible count distribution commonly used for biological count data is the negative binomial distribution. When researchers discuss zero-inflation, they refer to observation of more zeros than are expected by a particular distribution, such as the negative binomial distribution. The question is, does the negative control data contain more zeros than would be expected from a negative binomial distribution?

The negative binomial distribution has two parameters, a mean (or rate) parameter *μ* and an overdispersion parameter *ϕ*. Historically the negative binomial distribution has been used to model data where there is unknown random variation in the exposure, compared to a Poisson distribution (McCullagh and Nelder, 1989). The *ϕ* parameter can be seen as quantifying this unobserved variation. It can also be interpreted as biological variation on top of the technical variation due to sampling (Robinson et al., 2010), but this is not applicable for these negative control experiments.

For negative binomial distributions, two relations between summary statistics are particularly relevant for the question at hand. First, given *μ* and *ϕ*, the variance follows

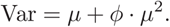

Second, given *μ* and *ϕ*, the probability of observing a count of 0 is

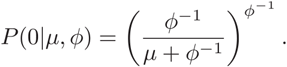

The overdispersion parameter *ϕ* can be assumed to be shared for an entire dataset, since it is a technical parameter when there is no biological variation. Each gene will have an independent mean *μ* within a dataset. Investigating relation between empirical mean and variance for each gene within a dataset illustrates a clear quadratic relation, with the exception of the Drop-seq data (Figure 1). From this relation a per-dataset *ϕ* parameter is fitted using least-squares.

**Figure 1.**
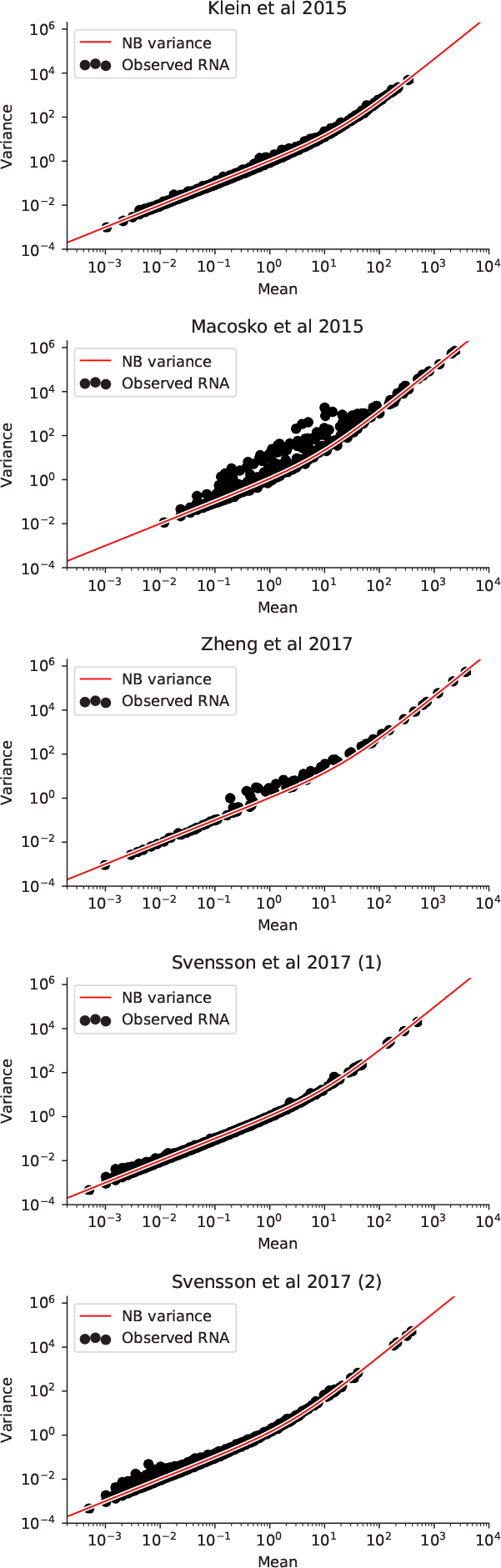
Mean and variance for negative control datasets. For each gene in each dataset the empirical mean and variance are calculated (black dots). The overdispersion parameter for quadratic trend is fitted for each dataset (red line).

Next the “dropout-rate” for each gene in each dataset is investigated. The dropout-rate is defined as the fraction of observed droplets (or cells) in which the gene has a zero value. This empirical measure can be compared to the expected fraction of zero-observations by calculating the probability of a count zero for the given mean and fitted overdispersion. For all genes in all datasets, except Drop-seq, the observation matches the expectation perfectly (Figure 2, Table 1).

**Figure 2.**
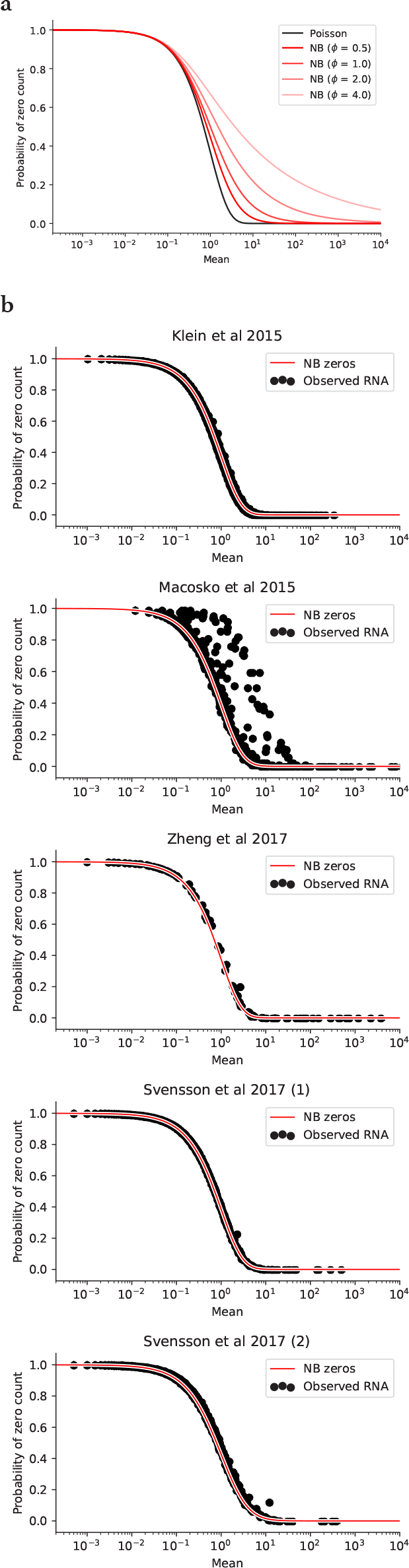
Expected and observed zeros. **a)** Theoretical fractions of zeros. Increasing the value of the overdispersion parameter will produce more zeros at higher mean expression levels. **b)** Observed dropout rates for every gene in each dataset given the mean (black dots), in relation to the expected number of zeros with a per-dataset overdispersion value and the mean of the gene’s expression level (red line).

## Discussion

The issue of zero-inflation and dropouts in scRNA-seq data has been discussed in the literature since the earliest scRNA-seq studies, and continues to be a topic of concern. While not recorded in literature, various personal interchanges indicates that it is a particular concern among researchers choosing between high-throughput droplet based methods and lower-throughput plate based methods.

Investigating negative control data for droplet based methods indicates the number of zeros observed is consistent with what is expected from count data. Additional zeros in biological data are likely due to biological variation.

Unfortunately no comparable negative control data exists for plate based methods. In a study about simulation of scRNA-seq data the investigators found droplet based data to be sufficiently modeled using negative binomial distribution, while plate based data needed zero inflation to be accurately simulated (Vieth et al., 2017). It is possible that UMI’s in droplet based methods deflates outliers in PCR duplicated counts. Another possibility is uneven sampling of fragments from gene bodies in plate based methods introduce an additional layer of count noise which give rise to gene-and-cell-specific overdispersion in addition to the global overdispersion, manifesting as additional zeros.

There is no clear record in the literature where the idea of zero-inflation in droplet based data came from. The conceptual origin seem to have been the combination of the zero-inflation model in SCDE attempting to model read counts in scRNA-seq data as well as observed bimodality on the log(TPM + 1) scale in a number of papers (Kharchenko et al., 2014; Shalek et al., 2013, 2014).

The distributions used here are taken as negative binomial, with per-dataset overdispersion, rather than Poisson. It is clear from the data in these negative control data that the total number of counts observed in each droplet has a large variation within a dataset, potentially giving rise to a negative binomial distribution from an underlying Poisson noise model^1^. It is not immediately obvious why this is since the starting material in every droplet will have the same volume and concentration. One possibility could be that beads used to deliver oligonucleotides into droplets have variable coverage of cDNAs, giving rise to variation in RNA capture rate per droplet.

Nevertheless, statistical analysis of data where noise follows the negative binomial distribution is simpler than analysis of a zero-inflated distribution. And the number of zeros observed can be decreased by counting more molecules through global increases in capture efficiency or increased sequencing depth per droplet.

## Methods

Count matrices for Drop-seq and Chromium data were generated as previously described (Svensson et al., 2017). The count matrix for the GemCode data was downloaded from the website linked to in the original publication (Zheng et al., 2017). The count matrix for the InDrops data was downloaded from GEO with accession GSE65525.

For ease of access, a folder containing all five datasets in H5AD format (Wolf et al., 2018) has been deposited to figshare^2^.

The per-dataset parameter was fitted using the `curve_fit` function in the `scipy. optimize` package.

All analysis and figure generation functions are available in a notebook at the same figshare project as the data, as well as on Github^3^.

## Footnotes

1 A previous version of this analysis was published at https://web.archive.org/web/20171119115058/http://www.nxn.se/valent/2017/11/16/droplet-scrna-seq-is-not-zero-inflated

1 See https://web.archive.org/web/20180225155427/http://www.nxn.se/valent/2018/1/30/count-depth-variation-makes-poisson-scrna-seq-data-negative-binomial/

2 Datasets analysed are avaiable at https://figshare.com/projects/Zero_inflation_in_negative_control_data/61292

3 Code for analysis reproduction is available at https://github.com/vals/Blog/tree/master/171116-zero-inflation

